# Gut^3^Gel as an in vitro model to investigate dietary modulation of the intestinal microbiota: An inulin supplementation case study

**DOI:** 10.1101/2025.05.29.656766

**Authors:** Miguel Antunes, Lucy Jongen, Abdul Mateen, João Sobral, Sebastião van Uden, Natalia Suárez Vargas, Daniela Pacheco

**Affiliations:** Bac^3^Gel, Lda, Taguspark - Innovation Building II, Av. Jacques Delors 411, 2740-122 Porto Salvo, Portugal

**Keywords:** Mucus-mimetic model, High throughput screening platform, preclinical testing, prebiotics, in vitro model, inulin supplementation

## Abstract

The intestinal microbiota plays a key role in human health, influencing digestion, immunity, and metabolism. While factors such as genetics and medications shape its composition, diet remains a primary driver of microbial modulation. Despite growing interest in using dietary interventions to beneficially alter the intestinal microbiota, assessing their effects in humans is challenging due to individual variability and the complexity of *in vivo* systems. This study explores Gut^3^Gel gradient colonic (G3GG) preclinical model as a tool to assess its representativeness on studying the effect of inulin supplementation on the intestinal microbiota of five healthy individuals compared to clinical evaluation studies. Significant inter-individual variability in baseline microbiota composition was observed, which strongly influenced microbiota response to inulin. While inulin supplementation led to a general decrease in alpha diversity, it significantly increased the abundance of health-associated genera such as *Bifidobacterium* and *Lacticaseibacillus*, along with enhanced microbial metabolic activity. Despite the intrinsic selectivity of G3GG for beneficial microbes, the model successfully captured inulin’s prebiotic effects and inter-individual differences, underscoring its relevance as a physiologically relevant high throughput tool for evaluating microbiota-targeting compounds and supporting personalized nutrition strategies.

## 1. Introduction

The intestinal microbiota plays a crucial role in human health and disease, influencing various physiological processes such as digestion, immune function, and metabolism^1^. The diversity and composition of the intestinal microbiota are shaped by multiple factors, including genetics, age, medication use (such as antibiotics), and environmental exposures^2^. However, diet stands out as one of the most significant determinants of microbial modulation^3^. Given this strong relationship between diet and microbiota, there has been increasing interest over the years in developing dietary interventions to positively modulate intestinal microbial composition^4^. One such approach is the consumption of prebiotics, which currently are broadly defined as substrates selectively used by host microorganisms that result in a health benefit^5^. The most well-known prebiotics include fructans (such as inulin) and galactooligosaccharides, both of which stimulate the growth of beneficial bacteria like *Bifidobacterium* and *Lactobacillus*^6,7^.

Despite the potential of prebiotics to modulate gut microbiota composition, studying their effects remains challenging. Human clinical trials, while necessary for demonstrating health benefits and efficacy of prebiotics, are often hindered by significant individual variability in microbiome composition and the complexity of *in vivo* environments, making it difficult to isolate the specific impact of dietary interventions^8– 10^. Contrasting, preclinical tools, such as *in vitro* intestinal models, provide a controlled setting that eliminates many external confounders and minimizes animal testing^11^. However, these models introduce their own set of limitations. For instance, the limited throughput of many *in vitro* systems reduces their ability to capture not only the inter-individual variability^12^, but also the initial diversity of intestinal microbiota impacted by host genetics, age, drug intake and epigenetics. Additionally, cost, technical, and operational limitations, as well as the requirement for specialized equipment, hinder their adoption^12^; these solutions are typically offered as services. The recently developed *in vitro* model, Gut^3^Gel gradient colonic (G3GG), offers a user-friendly, highly scalable, and cost-effective solution that overcomes existing limitations of *in vitro* models^13^. By modeling the physiological characteristics of intestinal mucus, G3GG serves as a promising tool for evaluating microbiome-modulating molecules such as prebiotics^13^.

In this study, the G3GG *in vitro* model was used to validate its potential as a testing platform for assessing the effects of prebiotics on intestinal microbiota composition. To achieve this, the modulatory effects of the well-characterized prebiotic inulin (derived from chicory roots)^10,14^ were monitored on the intestinal microbiota of five individuals. The impact of inulin was evaluated by analyzing changes in bacterial composition and metabolic activity. These analyses aimed to determine the ability of the G3GG model to simulate the intestinal mucus environment and serve as a robust tool for studying dietary interventions targeting intestinal microbiota.

## 2. Results

### 2.1. Inter-individual differences of intestinal microbiota samples have a substantial impact in assessing inulin prebiotic effect

The baseline microbial composition from the intestinal microbiota of five individuals (donors) was assessed to evaluate inter-individual differences. The samples used to assess the initial microbial composition (before inulin supplementation) correspond to 24 hours of stabilization of intestinal microbiota inocula in G3GG (Figure 1a). Even though, in general, the donor samples have similar microbial compositions, some variations can be observed. For instance, compared to the other donors, Donors 4 and 5 have substantially lower *Enterococcaceae* and Donors 2 and 5 have a high abundance of *Clostridiaceae*. Additionally, Donor 2 has an extremely low abundance of *Bifidobacteriaceae*, whereas Donor 4 has a substantially higher abundance compared to other donor samples, which is also observed for *Lactobacillaceae*. Furthermore, Donors 2 and 5 have a high abundance of *Enterobacteriaceae*. Comparison of baseline microbial compositions among donors, using unconstrained beta analysis, reveals that the donor-to-donor variability is a main driver of microbiome variation (PERMANOVA: R^2^ = 0.82; p-value = 0.001), which might overshadow the effects of inulin (Figure 1b). These results suggest that if there is an effect derived from inulin supplementation, it is likely only significant in specific donors or for specific bacteria.

**Figure 1.**
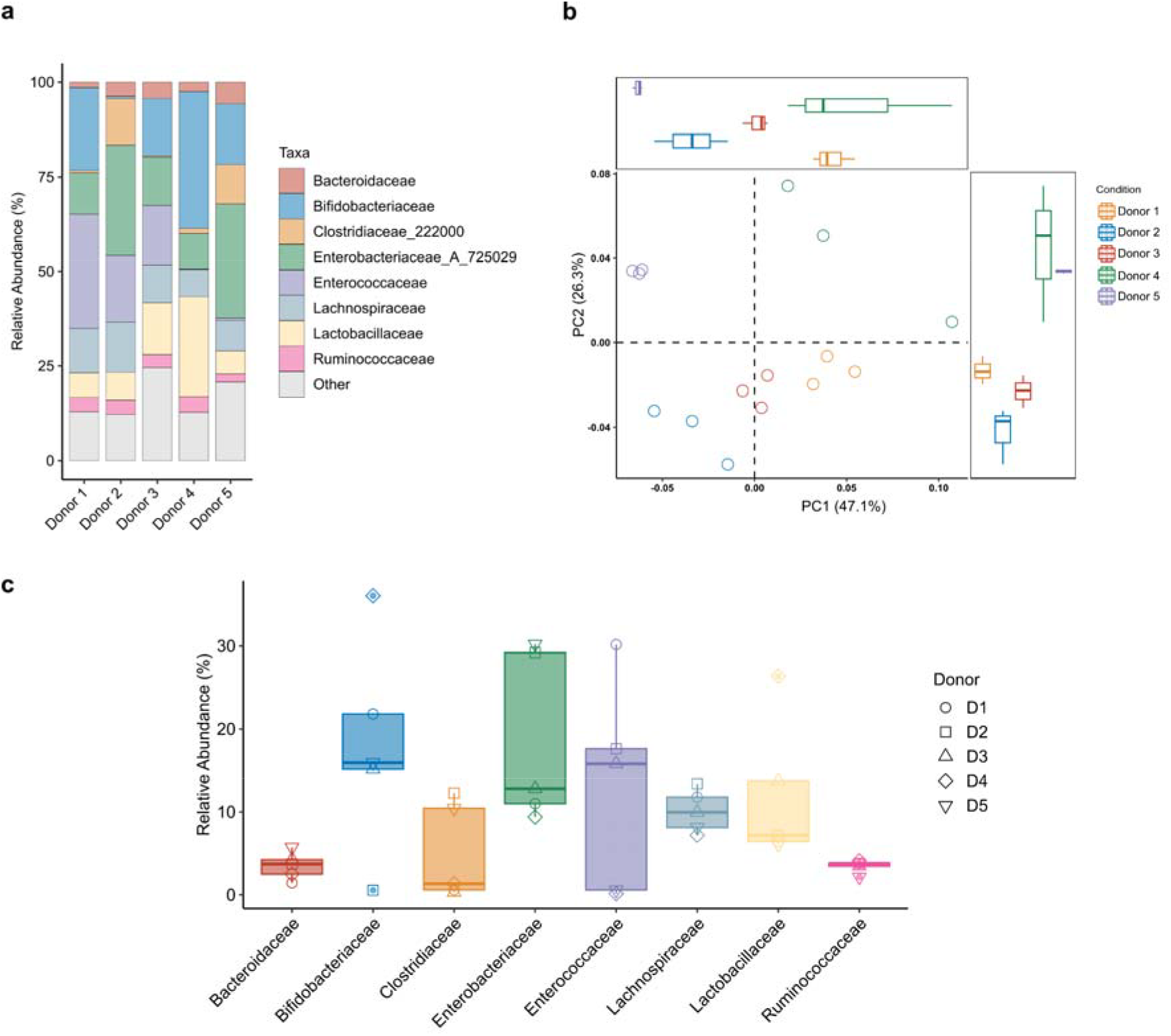
Inter-individual differences in baseline microbial composition are a dominant factor of variance. (**a**) Baseline microbial composition (following 24 hours of stabilization in G3GG; timepoint 0 hours) at the Family level presented as CLR transformed relative abundances (%) for each donor sample. (**b**) Beta diversity analysis determined based on Weighted UniFrac distance metric to evaluate dissimilarities in taxonomic composition between baseline microbial communities of the intestinal microbiota samples. Statistical significance was assessed using PERMANOVA. Side boxplots display the distribution of the samples along each principal coordinate axis, providing an overview of culturing condition dispersions. **(c)** Boxplots displaying the variability of the baseline microbial compositions at the Family level between donor samples presented as CLR-transformed relative abundances (%).

### 2.2. Microbial composition is shaped by inulin and incubation time despite large inter-individual variance

To investigate the suitability of G3GG as a preclinical tool for studying the effects of microbiota-modulating compounds, fecal samples were inoculated into G3GG with or without inulin and incubated for 48 hours. Samples were then collected for analysis of microbial composition and metabolic activity (Figure 2a).

**Figure 2.**
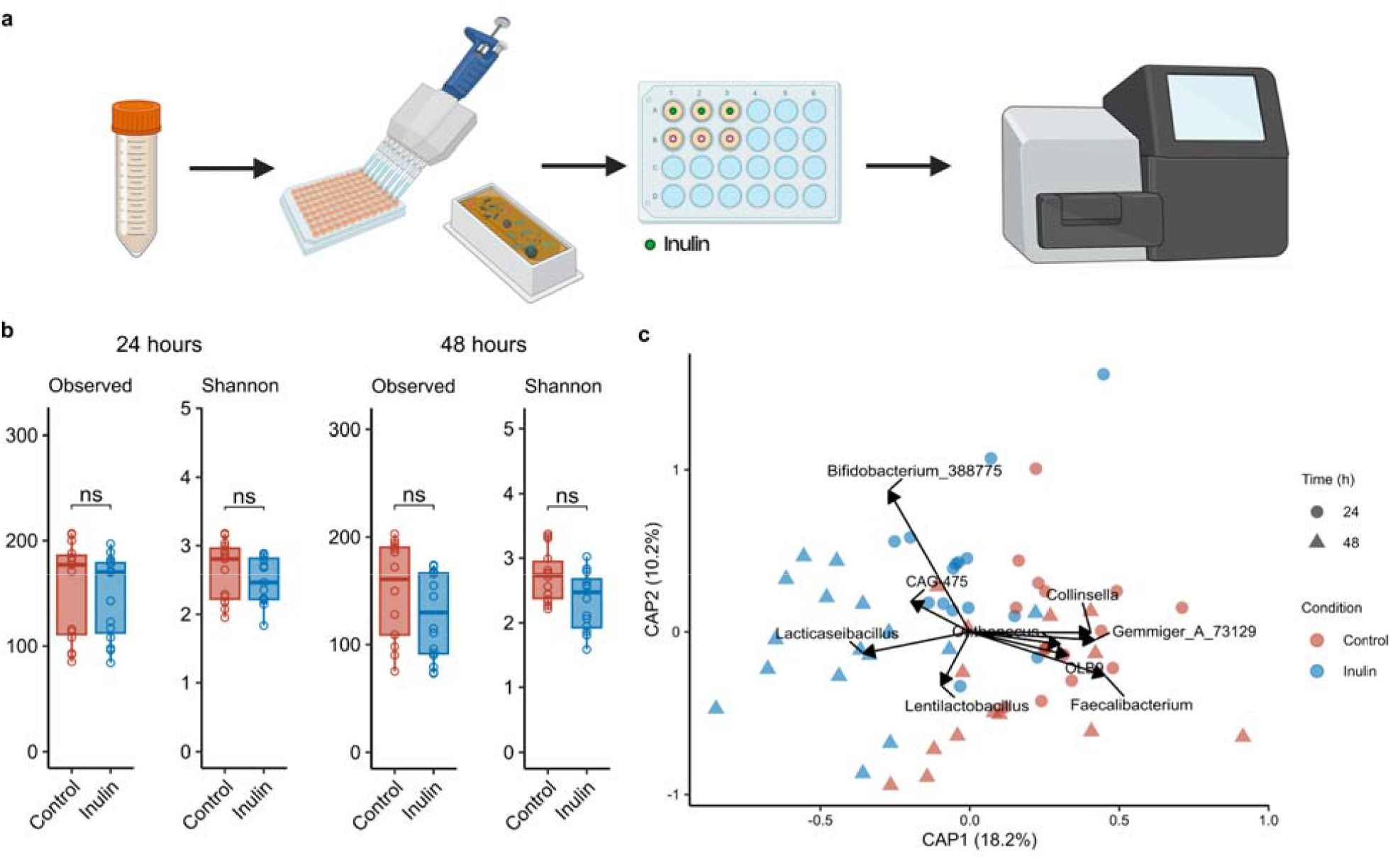
Inulin supplementation induces changes in microbial diversity and microbial composition in G3GG. (**a**) Schematic representation of inulin supplementation experiment. (**b**) Microbial diversity changes in response to inulin supplementation were investigated using alpha-diversity metrics: observed number of ASVs (richness) and Shannon diversity index (evenness) across donor samples at the incubation periods of 24 and 48 hours. Data normality was evaluated using the Shapiro-Wilk normality test (p-value < 0.05), and statistical significance was evaluated using the Wilcoxon rank-sum test (n = 15). (**c**) Constrained Principal Coordinates Analysis (CAP) based on Weighted UniFrac distance measure was performed on control and inulin supplemented samples across incubation time points (24 and 48 h) to determine the effect size of inulin supplementation. Inulin treatment and incubation time were used as constraining factors, with Donor set as a conditioning factor. Genera responsive to inulin supplementation were plotted on the ordination (Wilcoxon rank-sum test, p-value < 0.05).

To evaluate changes in alpha-diversity following inulin supplementation, the observed amplicon sequence variants (ASVs) and the Shannon index for the five intestinal microbiota samples were determined (Figure 2b). Incubation of intestinal microbiota with inulin for 24 or 48 hours resulted in a decrease in microbial diversity and evenness. However, this perceived decrease following inulin supplementation was not statistically significant.

Constrained Analysis of Principal Coordinates (CAP) using weighted UniFrac as the distance measure shows that, after controlling for inter-individual variability, the microbial composition is significantly affected separately by inulin supplementation (F = 11.27, p = 0.001) and incubation time (F = 9.15, p = 0.001). However, the impact of inulin supplementation does not seem to significantly depend on incubation time (F = 1.34, p = 0.251; Figure 2c). These results suggest that inulin supplementation and incubation time independently shape the microbiota composition, but the dominant factor influencing the microbial community structure remains the distinct baseline taxonomic profiles from donor samples, which account for 65.5% of the variation in microbial composition, whereas inulin and time together explained 10.2%.

### 2.3. Inulin supplementation significantly increases *Bifidobacterium* and *Lactobacillus* abundance

Inulin had a mild effect on taxonomic composition. At the Family taxonomic level, there was a general increase in *Bifidobacteriaceae* and *Lactobacillaceae*. Additionally, a decrease in *Enterobacteriaceae* was observed in donors following inulin supplementation.

The microbial selectivity of inulin was further evaluated at a higher taxonomic resolution by comparing the centered-log ratio (clr)-transformed abundances between the control and treatment conditions at the Genus level (Figure 3a). This analysis was complemented with a differential abundance analysis to identify significant alterations (Figure 3b). Inulin supplementation led to a significant increase in *Lacticaseibacillus* in all donor samples, particularly at 48 hours of incubation, as well as a significant increase in *Bifidobacterium, Ligilactobacillus* and *Lactobacillus* and a significant decrease in *Enterococcus*. Additionally, *Pedioccoccus* and *Levilactobacillus* increased in abundance in specific donors, whereas genera such as *Lentilactobacillus* and *Coprococcus* appear to decrease. The bifidogenic effect was observed for Donors 3 and 5, while lactobacilli increased significantly in abundance for Donors 1 and 2 (Figure 3c).

**Figure 3.**
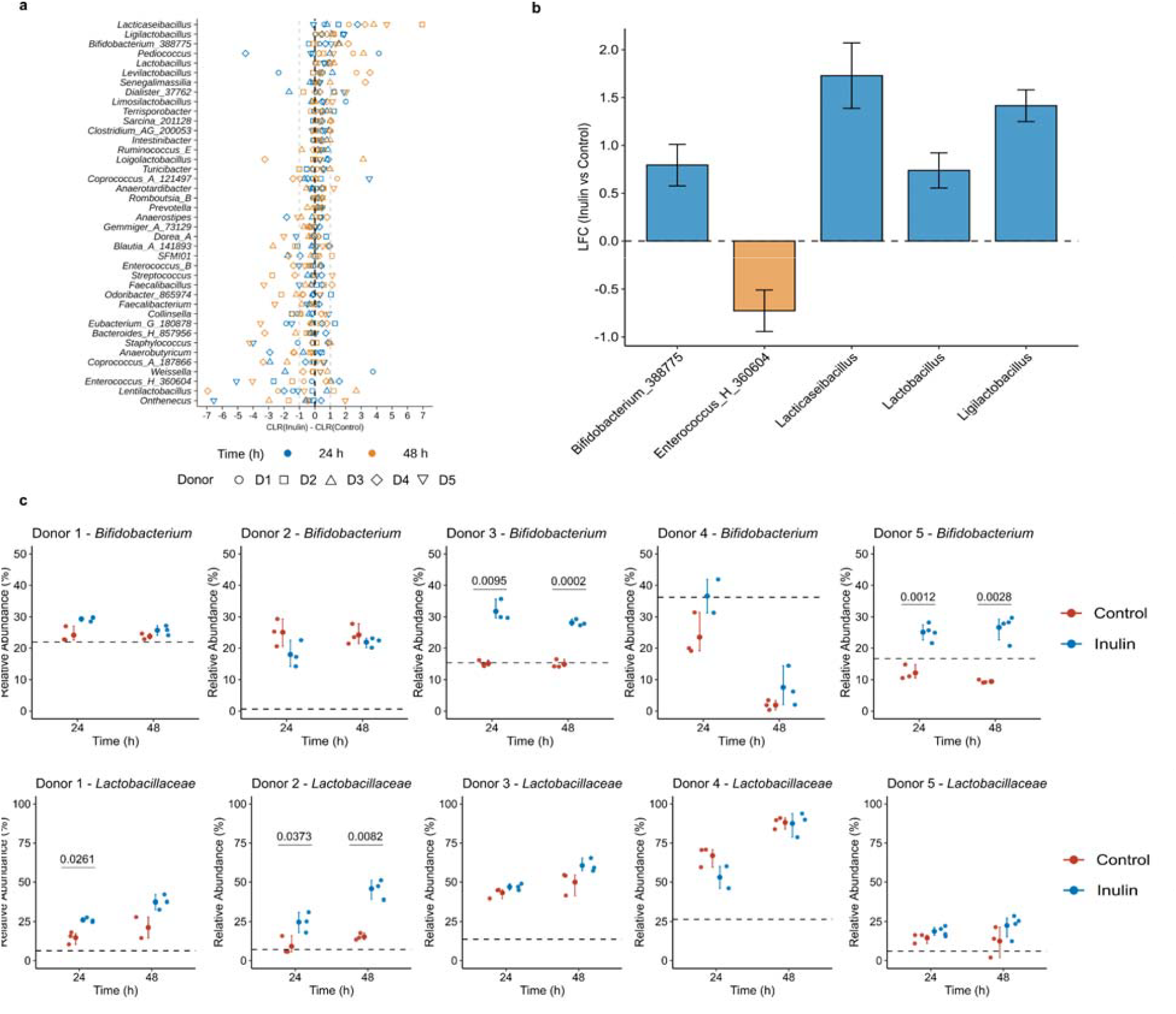
Inulin supplementation induces alterations on microbial composition. (**a**) Dot plot displaying the difference between CLR-transformed relative abundances of taxa with a prevalence above 10% and a mean relative abundance above the dataset median in G3GG over an incubation period of 48 hours. (**b**) Differential abundance analysis using ANCOM-BC2, accounting for donor variability across all donor samples and over an incubation period of 48 hours. The bar plot displays log fold change (LFC) values comparing inulin supplementation to control, with error bars representing standard error (SE). Multiple testing correction was applied using the False Discovery Rate (FDR) method with a significance threshold of 0.05. (**c**) Dot plots comparing the relative abundance (%) of *Bifidobacterium* and *Lactobacillaceae* between control and inulin-treated samples across donors and incubation times. Dashed lines indicate baseline relative abundances. Statistical significance was assessed using an independent t-test (n = 3), with error bars representing 95% confidence intervals.

### 2.4. Supplementation of Inulin results in an increase in microbial metabolic activity

Alterations in microbial activity within G3GG in the absence or presence of inulin were assessed using the MTT assay. Results revealed that microbial activity tended to decrease over the incubation period and that inulin supplementation led to increased microbial activity for Donors 1, 2 and 4, with a significant increase observed for Donor 2 (Figure 4a). To assess global microbial activity changes upon inulin supplementation while accounting for inter-individual variability, a linear mixed-effects model (LMM) was applied for each time point (24 and 48 hours). The analysis revealed that inulin supplementation results in a significant increase in microbial activity compared to the control at 48 hours (β = 0.52, SE = 0.20, p = 0.016), but not at 24 hours (β = 0.11, SE = 0.09, p = 0.259). These results suggest that inulin supplementation is enhancing microbial activity independently of donor-specific differences in microbiota composition.

**Figure 4.**
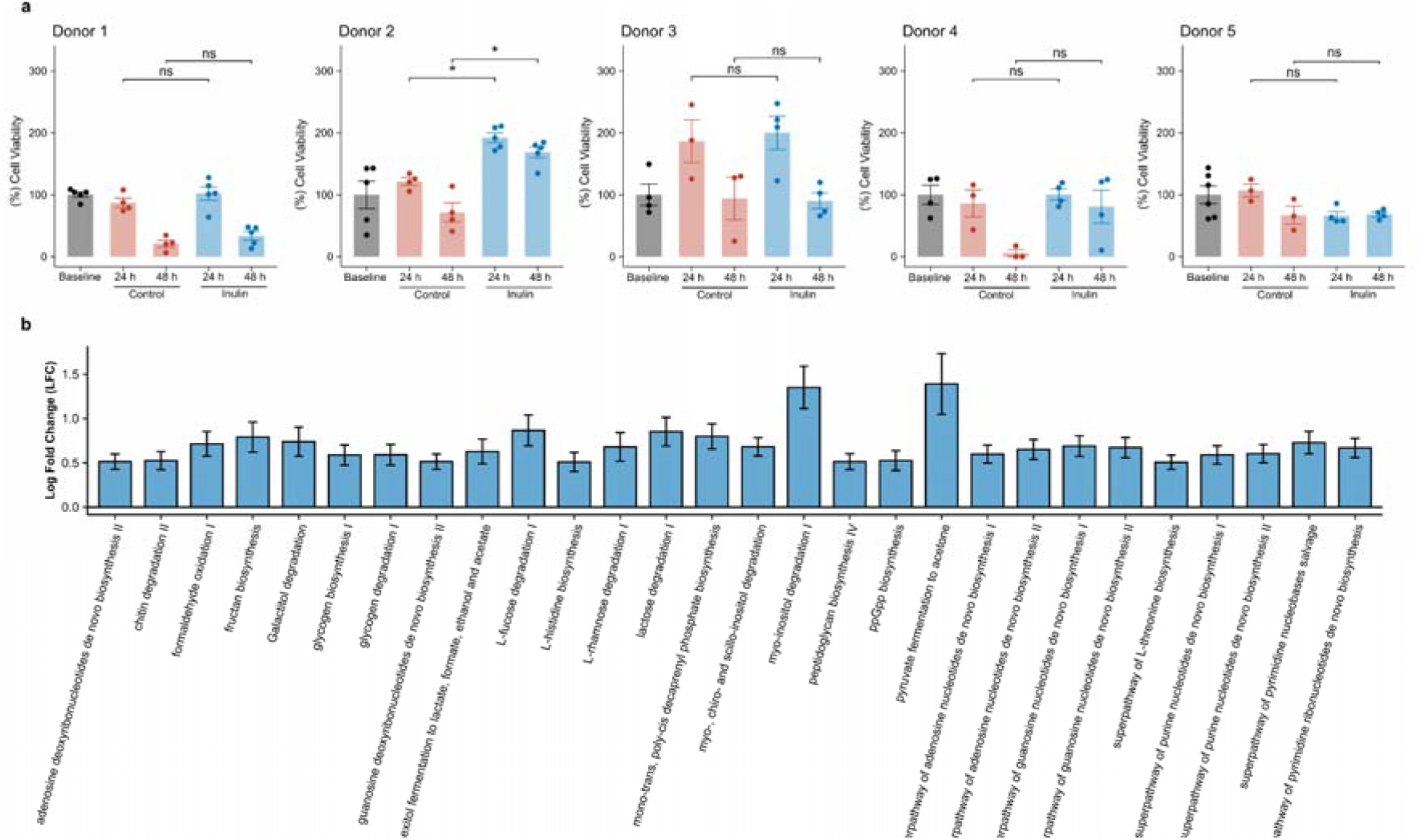
Inulin supplementation affects microbial activity. **(a)** Bar plots displaying the percentage of metabolically active cells after 24 or 48 hours of incubation with inulin. Metabolic activity is expressed relative to baseline samples (0 hours, prior to inulin supplementation), which were set to 100%. Error bars represent the standard error of the mean (SE). Statistical significance was determined using the Wilcoxon rank-sum test, with a minimum of four technical replicates per condition. **(b)** Predictive functional profiling using PICRUSt2. The bar plot shows MetaCyc pathways with statistically significant differences in log fold change (LFC) between inulin supplementation and control conditions. Only pathways with an absolute LFC greater than 0.5 (LFC > 0.5 or < -0.5) are displayed. Error bars represent the standard error of the mean (SE).

The metabolic pathways enriched upon inulin supplementation were inferred using PICRUSt2. Notably, several carbohydrate metabolism pathways showed increased predicted abundance, including myo-inositol degradation I, lactose degradation I, myo-, chiro-, and scyllo-inositol degradation, L-fucose and L-rhamnose degradation I, and glycogen metabolism. Additionally, there was an increase in the pyruvate fermentation to acetone, hexitol fermentation to lactate, formate, ethanol, and acetate pathways, suggesting the activation of short-chain fatty acid (SCFA) production pathways and fermentation activity associated with inulin prebiotic effects. Several amino acid biosynthesis pathways, as well as nucleotide salvage were also significantly enriched.

## 3. Discussion

This study investigated the potential of G3GG as a platform to assess the effects of inulin, a highly characterized prebiotic compound both *in vitro* and *in vivo*, on intestinal microbiota composition and metabolic activity.

Although not statistically significant, a notable reduction in alpha diversity following inulin supplementation was observed, which is consistent with previous randomized clinical trials^15^. This decrease in diversity is likely associated with the observed significant increase in the relative abundance of specific bacterial taxa, particularly *Bifidobacterium* and *Lactobacillus*, whose growth has been previously shown to be stimulated by fructo-oligosaccharides and inulin supplementation^6,7,16,17^.

Inter-individual variation was identified in this study as a significant factor influencing the assessment of general trends from prebiotic interventions. Previous studies have shown that consumption of prebiotics to target specific microbes conferring health benefits is only successful in a subset of respondent individuals likely due to highly variable baseline microbial compositions^18–21^. Differences in microbial community composition among the five donors’ samples before supplementation with inulin were evident. Concomitantly, culturing in G3GG led to an intrinsic prebiotic effect by promoting the growth of bacteria such as *Bifidobacteriaceae* and *Lactobacillaceae*, as previously shown^13^. While this selectivity of G3GG may partially mask the effects of the test compounds on microbial composition, it may also present an advantage as it provides a growth environment that mimics the native intestinal mucus layers. This contrasts with traditional *in vitro* models that lack a mucus component, which may not fully capture microbial dynamics occurring at the mucosal surface. G3GG provides a physiologically relevant context that can be customized with more preferable mucin types, and that can be exploited to evaluate the response to microbiota modulation interventions. Despite the influence of G3GG on *Bifidobacterium*, and *Lactobacillus*, the effect of inulin on these bacterial genera was effectively detected.

Inulin induced pronounced alterations compared to the control condition. Inulin supplementation led to a significant increase in *Bifidobacterium* and *Lacticaseibacillus* accompanied by significant increase in metabolic activity at 48 hours of incubation. These results are consistent with inulin being described as a rapidly fermented fiber, typically leading to quick and pronounced changes in microbial composition and particularly promoting the growth of *Bifidobacterium* and *Lactobacillus* species^22^. Furthermore, a significant decrease in *Enterococcus* abundance with inulin was observed in this study. Although this is not a commonly reported effect of inulin, it has been observed in a few studies^23–25^.

Associations between the baseline abundance of *Bifidobacterium* and the bifidogenic effect of inulin have been reported in previous studies. In particular, individuals with low baseline levels of *Bifidobacterium* tend to exhibit a greater relative increase in response to inulin supplementation^26^. However, in this study, the greatest increase in *Bifidobacterium* was observed in donor samples with intermediate baseline abundances (15-25%). In contrast, donors with either very high (Donor 7) or very low (Donor 4) baseline levels did not show substantial changes following inulin treatment. This apparent discrepancy may be attributed to the small sample size used or to unaccounted confounding factors. One possible explanation is variation in habitual dietary fiber intake, which has been shown to modulate the responsiveness of microbiota to prebiotic interventions^16,27^. As this variable was not controlled for in the present study, it may have influenced individual responses to inulin.

In summary, G3GG is shown to be a suitable platform for testing the efficacy of compounds potentially conferring health benefits, particularly inulin. This model effectively captures inter-individual variability in a high throughput manner. G3GG is a cost- and time-efficient tool to determine the efficacy of microbiome-modulating compounds and may also contribute to personalized dietary interventions.

## 4. Materials and Methods

### 4.1. Donor recruitment

All donors provided written informed consent, confirming their understanding of the study and their voluntary participation. Approval from an ethical committee was not required for this study, as it did not involve prospective evaluation or the use of laboratory animals. The study was conducted exclusively employing non-invasive methods, specifically the collection of fecal samples. Furthermore, all donors’ personal information was kept anonymous upon sample collection and throughout data analysis.

### 4.2. Isolation of intestinal microbiota

Intestinal microbiota was isolated from fecal samples from five healthy human adults following the European Guidelines of Fecal Matter Transplantation^28^ and as previously described^13^. Briefly, a minimum amount of 30 g of fecal sample was suspended in 150 mL of 0.9% saline solution using a blender, followed by a filtration step. Furthermore, 45 mL of the filtrate (stool solution) was transferred to 50 mL falcons with 5 mL of 100% glycerol and stored at -80°C until needed. All donors were European, aged between 25 to 36 years, with a BMI ranging from 18.5 to 30, without gastrointestinal complaints or any related disorders, and free of antibiotics or food supplements at least 3 months prior to this study.

### 4.3. Test compounds

The test compound used was inulin from chicory roots (Sigma Aldrich, CAS 9005-80-5). An inulin stock solution was prepared by dissolving inulin in 0.9% NaCl. These solutions were then heated to 30°C to ensure a complete dissolution, filter sterilized (0.22 μm filters) and stored at 4°C.

### 4.4. Gut^3^Gel gradient colonic (G3GG)

Gut^3^Gel gradient colonic (G3GG colonic, Bac^3^Gel Lda) is a commercially available hydrogel-based structure mostly containing 50 mg.mL^−1^ porcine stomach type III mucin (Sigma-Aldrich, M1778; Germany); previously reported concentration in intestinal mucus^29^, produced in Brain Heart Infusion broth medium (BHI, Sigma-Aldrich)^13^. G3GG mimics several key characteristics of the gastrointestinal tract mucus such as chemical composition, viscoelastic properties, and a gradient cross-linking pattern with intercalated mucins in its structure with a three-dimensional gradient of oxygen tension^13^. The method of production is patented (WO2020128965A1)^30^.

### 4.5. Inulin treatment of microbiota in G3GG

Five Intestinal microbiota samples, stored at -80°C, were thawed in a water bath set at 37°C and diluted with BHI broth medium in 1:1 ratio. Then, 700 μL of the diluted microbiota from each sample was inoculated, in triplicate, in G3GG and incubated at 37°C for 24 hours for the stabilization of the microbial communities. Following the stabilization period, the supernatant on the surface of G3GG samples was discarded. Stabilized microbiota in G3GG was then supplemented with 700 μL of inulin stock solution (treatment group) so that the final concentration was 5 g.L^−1^. For the control group, 0.9% NaCl solution was added instead of the inulin solution. The samples were then incubated at 37°C for 24 and 48 hours. Sterile G3GG was included as a control for sterility, rheological properties, oxygen tension, and sequencing.

### 4.6. Analysis of microbial community composition

For each time-point, the supernatant of G3GG was discarded and G3GG was washed with 0.9% NaCl solution, so that only the microbes that colonized the G3GG structure were considered. To retrieve the bacteria within G3GG, 50 mM of sodium citrate (Sigma) was added (dilution factor = 1.5) to fully dissolve the G3GG structure. The dissolved gel was transferred into 2 mL microtubes and centrifuged (16000 x g) for 10 minutes. The pellet was resuspended with 800 μL of the CD1 solution from the QIAamp Power Fecal Pro DNA Kit and processed following the protocol provided by the manufacturer.

To assess microbial community composition, the V3-V4 hypervariable regions of the 16S ribosomal RNA (rRNA) gene were targeted for bacterial identification, using primer pairs referenced by Klindworth et al. and Herlemann et al.^31,32^, and following the 16S Metagenomic Sequencing Library Preparation Illumina protocol (Part # 15044223 Rev. B, Illumina, CA, USA). The ZymoBIOMICS Microbial Community Standard (Catalog D6306, Zymo Research, Irvine, CA, USA) was utilized as a quality control assessment to evaluate potential biases and artifacts in the PCR and sequencing steps. Library preparation and amplicon sequencing (Illumina Miseq, 2×300 cycles) were performed at the NGS laboratory at Bac^3^Gel, Portugal.

The raw paired-end reads were primer-trimmed using cutadapt. Analysis of the 16S amplicon sequences was performed using QIIME2 (version 2024.10) and R (v4.4.2) in RStudio (version 2024.12.1). Denoising and generation of amplicon sequence variants (ASVs) was performed using the DADA2 plugin within QIIME2. Taxonomic assignment of ASVs were determined with the q2 feature-classifier plugin against the Greengenes database version 2024.09 using a classifier trained on the V3-V4 regions. Low-abundance ASVs that were present with a total ASV count of less than 0.1% the maximum count was excluded. Furthermore, ASVs that were taxonomically unclassified at phylum rank or taxonomically assigned to mitochondria or chloroplast were removed from the abundance table.

Alpha diversity analysis (Shannon diversity index) was performed using the R package phyloseq v1.46.0. Beta diversity analysis and ordination plots were created using the R packages phyloseq and ape v5.8. Principal coordinate analysis (PCoA) was employed as the ordination method, with weighted UniFrac used as the distance measure. For compositional data analysis, zero values in the abundance table were replaced using the count zero multiplicative method with the R package zCompositions v1.5.0.3. The data was then transformed using the centered log-ratio (CLR) method. Exploratory data analysis was conducted through compositional principal component analysis (PCA) of the datasets, achieved via singular value decomposition (SVD) of the CLR-transformed data. Constrained analysis of principal coordinates (CAP) with Weighted UniFrac as distance metric was used to determine the effect of inulin on the microbial composition. Inulin and incubation time were set as constraining factors and donor as a conditioning factor. Differential abundance analysis was performed using ANCOM-BC2 v2.10. Treatment, incubation time and donor were set as fixed effects. Samples with library sizes less than 100 were excluded from the analysis and multiple testing correction was applied using the False Discovery Rate (FDR) method with a significance threshold of 0.05. PICRUSt2 v2.6.2^33^ was used to predict the functional profiles of microbial communities based on the 16S rRNA gene sequences. Significantly enriched pathways were determined using ANCOM-BC2. To reduce the influence of extremely rare taxa and improve statistical robustness, features were previously filtered to retain only those present in at least 10% of samples and with a mean relative abundance above the dataset median.

### 4.7. Metabolic activity assay

To evaluate the metabolic activity of the microbial communities within G3GG either in the absence or presence of inulin, the Cell Proliferation Kit I (MTT) (Roche) was used. Briefly, for each time point, 100 μl of dissolved gel samples were mixed with 10 μl of MTT labelling reagent and incubated for 4 hours at 37°C. Following the incubation period, 100 μl of solubilization solution was added and samples were incubated overnight at 37°C. The optical density was measured at 570 nm with a reference wavelength of 670 nm using the SPECTRAmax plus 384 microplate spectrophotometer (Molecular Devices). Sterile G3GG without microbiota was used as the blank.

## Data availability statement

The data for this study have been deposited in the European Nucleotide Archive (ENA) at EMBL-EBI under accession number PRJEB89648 (https://www.ebi.ac.uk/ena/browser/view/PRJEB89648).

## Acknowledgments

The authors would like to thank the European Innovation Council through the Accelerator mechanism (HORIZON-EIC-2023-ACCELERATOROPEN-01 ID: 190135075) for partially funding the validation of the technology to be employed as intestinal models to accelerate drug development.

